# Many-body Interaction Competition Drives Reentrant Phase Transitions

**DOI:** 10.64898/2026.05.26.727925

**Authors:** Jiale Qiao, Rob M. Scrutton, Daoyuan Qian, Tuomas P. J. Knowles

## Abstract

Reentrant phase transitions, in which multicomponent systems phase separate at intermediate concentrations but dissolve at experimentally accessible higher concentrations, are ubiquitous in mixtures such as biomolecular condensates. We show that introducing reversible dimerization into a multicomponent Flory–Huggins model and integrating out the dimer state generate an effective three-body repulsion that reshapes phase diagram geometry. This emergent higher-order interaction arises naturally from an interaction competition and mass-action equilibrium, providing a microscopic explanation to the previously phenomenological three-body interaction. We derive a closed-form phase boundary equation capturing phase separation and reentrant dissolution in this minimal model, and predict explicit interdependence between competition strength, emergent many-body interactions, and dissociation constants. We recover and extend the reentrant phase boundary scaling relations through interaction renormalization, with regime of validity. We apply our model to G3BP1–RNA–suramin and explain the underlying mechanisms from the physical parameters inferred from reentrant phase boundaries.

## I. INTRODUCTION

Reentrant dissolution describes a multicomponent system that phase separates [1] at intermediate concentrations and reenters the homogeneous state upon further addition of one component [2, 3]. Such behavior is one of the most important physical processes in biology, playing a central role in cellular stress response [4], cellular regulation [5–7], and disease [8–10]. Reentrant dissolution has been increasingly observed in biomolecular condensates involving proteins, nucleic acids, salts, and small molecules [11, 12]. With high-throughput platforms [13], an increasing number of finely resolved phase diagrams have been measured [14, 15]. Despite the increasing number of experimental observations, the microscopic mechanisms of reentrant phase transitions are still not fully understood.

Reentrant dissolution in biomolecular systems is commonly attributed to competition between intermolecular forces that alternatively promote or suppress phase separation [16]. For example, the ribonucleoprotein (RNP)-RNA complex exhibits reentrant dissolution at high RNA concentration [17]. The behavior is explained by the interplay between sequence-encoded short-range electrostatic attraction and long-range Coulomb repulsion due to charge inversion in the complex. Reentrant dissolution arises when RNA concentration increases beyond charge neutrality and acquires repulsion. More recently, electrostatics alone have been shown to generate salt-dependent reentrant phase behavior [18]. In Caprin1 and its variants, multivalent counterions promote condensation by bridging protein chains and inducing effective attractions, while increasing salt suppresses phase separation through screening and ion partition entropy. Such interplay produces reentrant dissolution, highlighting that effective many-body interactions can arise from ion partitioning and electrostatic networks. Although these successful mechanisms remain system-dependent, phenomena such as electrostatic screening [14], charge inversion [17], and ion bridging [18] share a common underlying microscopic structure.

The sticker–spacer framework provides a conceptually simple and powerful description of associative polymer networks [12, 19, 20]. In this model, multivalent binding sites (stickers) are connected by flexible linkers (spacers), and phase behavior reflects the interplay between intermolecular sticker–sticker attraction and intramolecular spacer-mediated interactions. Reentrant dissolution can emerge naturally because the model captures both intermolecular interactions and condensate network connectivity. However, the increased microscopic complexity often necessitates numerical treatments [21, 22], limiting analytical tractability. Moreover, quantitative predictions typically require additional assumptions and estimates including interaction strength, binding affinity, and polymer architecture. Other theoretical frameworks, including the Voorn–Overbeek model [23], random-phase approximation [24, 25], and transfer-matrix model [26, 27], as well as newly developed computational Mpipi model [28] and machine learning model [29], provide complementary descriptions of polymer phase behavior with different levels of microscopic detail. However, they often rely on phenomenological parameters and additional assumptions for quantitative predictions.

Mean-field approaches to explain reentrant dissolution typically employ the multicomponent Flory–Huggins free energy [30, 31]. Numerical studies have shown that cubic contributions can substantially alter multicomponent phase behavior and generate complex phase diagrams by promoting or opposing phase separation [32]. However, higher-order coefficients are introduced phenomenologically, without a microscopic derivation based on binding processes in biological systems. Complementary to such free-energy interaction descriptions, alternative phenomenological framework has been proposed that encodes condensate formation in the solubility product [33]. Reentrant phase boundaries are derived analytically [34] from a mass-action equilibrium with variable stoichiometry and oligomers, although the underlying microscopic mechanisms are not explicitly resolved.

More recently, geometric tie-line analysis [35–37] has clarified how effectively repulsive many-body interactions shape the reentrant phase boundary geometry without relying on derivative-based coexistence conditions which often prevents analytical access. This approach reveals scaling relations [36] that describe phase separation and reentrant dissolution boundaries, but the effective higher-order interactions remain imposed parameters rather than emergent quantities derived from microscopic physics.

In summary, previous attempts to explain reentrant dissolution have focused on microscopic electrostatic mechanisms, molecular binding models, or higher-order free energy corrections. Although these approaches capture different aspects of the same phenomenon in different representations, how they connect to explain reentrant dissolution remains an open question.

Here we show that reentrant phase transitions can arise from an interaction competition. We develop a minimal quantitative theory by introducing reversible dimeriza-tion into a multicomponent Flory–Huggins model and integrating out the dimer state perturbatively. This renormalization procedure reveals an effective higher-order interaction originating from competition Δ*χ* defined in Eqs. (2)–(3), representing the interplay between monomer–monomer and monomer–dimer interaction channels coupled with mass-action equilibrium. Reentrant dissolution geometry requires that the competition is positive and therefore the three-body interaction is effectively repulsive, suppressing condensate network formation at higher concentrations in experiments. The resulting effective three-body interaction *χ*^(3)^ ∝ Δ*χ* × *K* is proportional to the competition Δ*χ* and the association constant *K*. This mechanism provides a microscopic interpretation for the higher-order interaction terms that have previously been introduced phenomenologically.

Using a tie-line formulation, we derive an analytical closed-form phase boundary equation that captures phase separation and reentrant dissolution governed by physically interpretable parameters. The equation shows that the established phase-boundary scaling relations [36] emerge as the low-concentration limit of the theory, while naturally extending to higher concentrations in experiments through an interaction corrections. The resulting theory predicts explicit connections between competition strength, association equilibrium, and reentrant boundary geometry. We apply our theory to the G3BP1-RNA-suramin system [38] to investigate the mechanisms underpinning the reentrant phase transition by extracting the key physical parameters from phase boundaries. We hence demonstrate that the observed reentrant behaviors are quantitatively consistent with this shared microscopic mechanism.

### II. THEORETICAL FRAMEWORK

The central concept of our model is the newly defined competition between two-body interaction channels. This competition arises naturally when dimerization from existing monomer components perturbs the two-component Flory–Huggins free energy

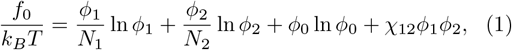

where *N*_1_, *N*_2_ are the polymer lengths in lattice unit *v*_0_; *ϕ*_1_, *ϕ*_2_ are the volume fractions of components 1 and 2. *ϕ*_0_ = 1 −*ϕ*_1_ − *ϕ*_2_ is the volume fraction of the solvent component 0. *χ*_12_ *<* 0 is the 1-2 interaction driving phase separation. Dimerization in the dilute phase captures the minimal condensate network formation and allows analytical treatment. The dimer state should be interpreted as the minimal coarse-grained object whose elimination generates effective interactions. The physical mechanism does not rely on the existence of stable thermodynamic dimers. As shown in Appendix A 1 and B, the competition is derived directly from integrating out dimer states. A closer examination shows that the interaction competition is the difference between the monomer–dimer and monomer–monomer interactions [Fig. 1(b)–(d)], weighted by a volume coefficient 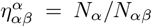 (the volume fraction of monomer *α* in dimer *αβ*). We assume the incompressibility condition *N*_12_ = *N*_1_ + *N*_2_ so that 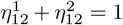. In terms of the two-body interaction *χ*_*αβ*_, the competition parameters take the form

**FIG. 1.**
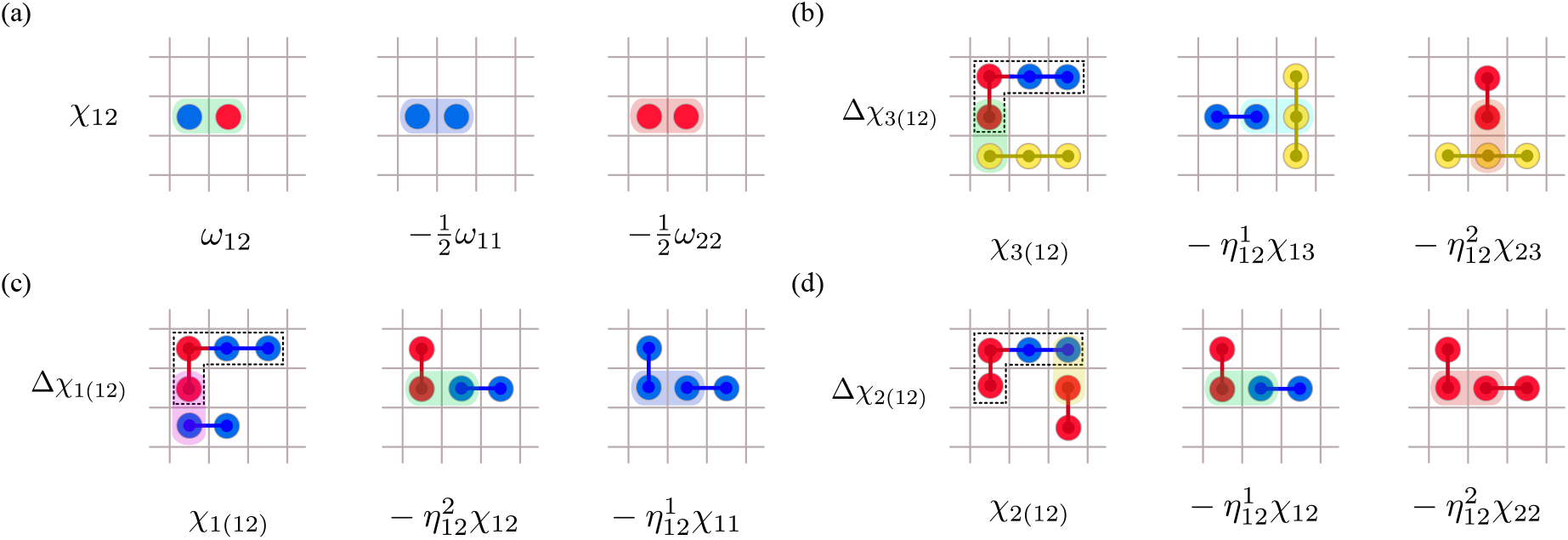
Schematic of the two-body interaction *χ*_12_ and competition Δ*χ*. (a) The two-body interaction parameter *χ*_12_ in standard Flory-Huggins model is interpreted as the contact energy difference when switching on heterogeneous interaction *ω*_12_. (b) The competition Δ*χ*_3(12)_ [Eq. (4)] is defined similarly as the energy difference in dimer (12)-monomer (3) channel and monomer (1,2)-monomer (3) channel with volume coefficients 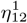 and 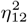. The interaction channel *χ*_3(12)_ is generally different from the channels *χ*_13_ and *χ*_23_. (c) Δ*χ*_1(12)_ is the self-competition limit with component 3 → 1, where *χ*_11_ = 0. (d) Δ*χ*_2(12)_ is the analogous case with 3 → 2 and *χ*_22_ = 0.

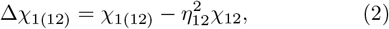

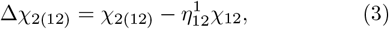

where we enclose the dimer index in parentheses, (12), to show explicitly its constituent monomers and to indicate that it is treated as a single independent species. In three-component system, the competition parameter reveals a more complete and general structure

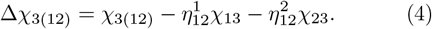

The definition of competition is directly analogous to the Flory–Huggins 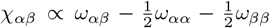 [Fig. 1(a)]. While *χ*_*αβ*_ measures the free energy perturbation upon switching on heterogeneous *αβ* interactions, competition Δ*χ* measures the free energy perturbation from dimer formation. Competition can be alternatively interpreted in terms of the molecular interaction energy Ω_*αβ*_ = *N*_*α*_*N*_*β*_*χ*_*αβ*_ which scales with the lengths *N*_*α*_ and *N*_*β*_ of the interacting molecules. In this representation, the molecular competitions are ΔΩ_1(12)_ = Ω_1(12)_ −Ω_12_ and ΔΩ_2(12)_ = −Ω_2(12)_ Ω_12_, and in the three-component case ΔΩ_3(12)_ = Ω_3(12)_ −Ω_13_ −Ω_23_.

The free energy after dimerization [Eq. (A2)] is minimized under the conservation of volume, which is also the number of molecules, constraint [Eq. (A3)]. The chemical potential before and after dimerization should be at equilibrium [Eq. (A6)]. In Eqs. (A7)–(A13) we identify an effective dimer association constant with

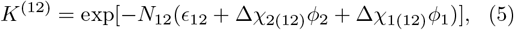

where *K*^(12)^ exhibits a weak exponential decay with *ϕ*_1_, *ϕ*_2_ set by Δ*χ*_1(12)_ and Δ*χ*_2(12)_. The intrinsic associa-tion constant 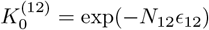 corresponds to zero competition Δ*χ* = 0. *ϵ*_12_ is the free energy change in the dimerization reaction. In Eq. (A14), we find that this competition modification gives correction in the same exponential form as the depletion correction by further solving mass-action equilibrium Eq. (A10) iteratively to higher order, which identifies the perturbative regime of validity:

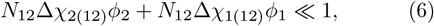

which is generally wider than the regime of validity for an exponential scaling [Fig. 2(b)] of reentrant boundary

**FIG. 2.**
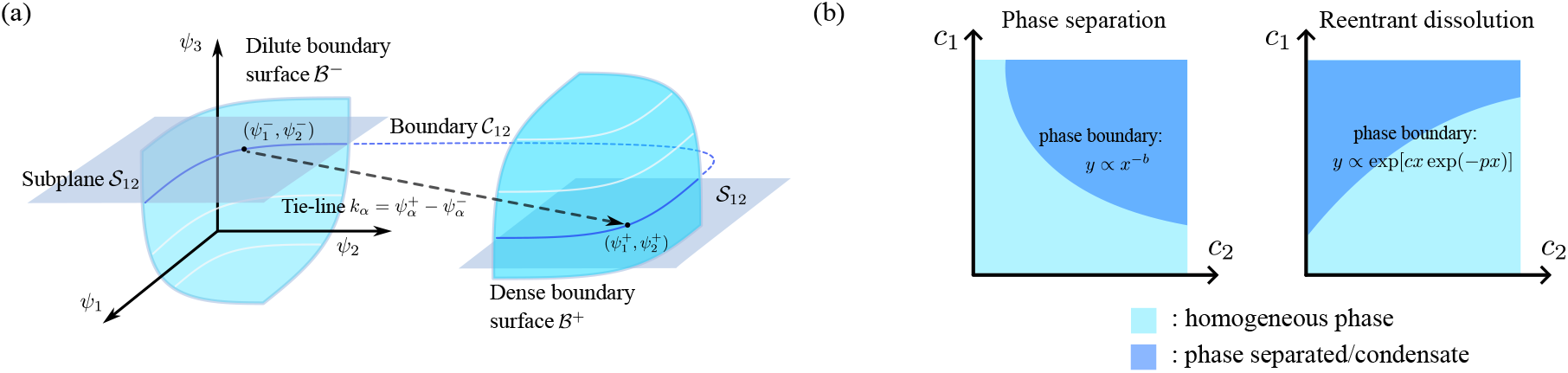
Geometry of multicomponent phase space and phase boundary. (a) The mixtures in experiment are generally multi-component. The phase boundary hypersurface *ℬ* encloses the phase-separated region in multicomponent composition space. The tie-line vector 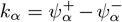 connects the coexisting composition 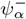 on the dilute branch *ℬ*^*−*^ and 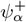 on the dense branch ℬ^+^ as determined by the common-tangent construction. The experimentally measured phase boundary *𝒞*_12_ in component 1 *–* 2 subspace is given by the intersection of the plane *𝒮*_12_ with the hypersurface *ℬ*. (b) The phase boundary model [Eq. (11)] implies the scaling relations adapted from Ref. [36]: *y* ∝ *x*^*−b*^ in phase-separated regime, and *y* ∝ exp[*cx* exp(*−px*)] in reentrant dissolution regime, with a correction suppressing term exp(*−px*).

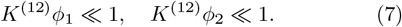

We then integrate out the dimer state under perturbative calculation, where the previous two-body terms generate the effective three-body repulsive terms in the interaction part of the free energy Eq. (A16) but leave the entropy part unchanged Eq. (A19). Collecting results, we show that the effective three-body interaction parameter is proportional to the competition and association constant, with a prefactor counting allowed interaction channels (see Appendix B for the general case)

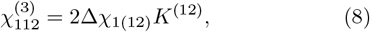

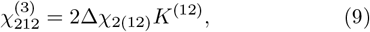

where we denote the concentration-independent part 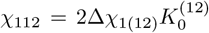 and 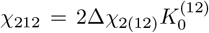. The renormalized free energy describing reentrant dissolution takes the form

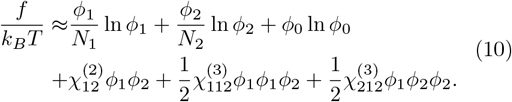

In this representation, the interaction coefficients carry weak *ϕ* dependence. It can be shown that, by expanding these coefficients to leading order Eq. (A23), we obtain a constant coefficient free energy [Eq. (A25)– (A28)] that effectively captures the same physics.

In the Flory–Huggins lattice framework, volume fraction and molar concentration are connected by *ψ*_*α*_ = *v*_0_*N*_*α*_*c*_*α*_, where *v*_0_ is the volume of 1 mol referencing molecules such as water. Note we use tie-line analysis conventional *ψ*^±^ to label phase diagram volume fractions instead of the generic free-energy variable *ϕ*.

Tie-line analysis describes the geometry of multicomponent phase space [Fig. 2(a)] where the tie-line vector 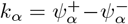 connecting two sides of coexistence boundaries from common tangent construction encodes thermodynamic information [35]. In terms of molar concentration, the phase boundary equation C_12_ in concentration subspace S_12_ derived in Appendix A 2 is

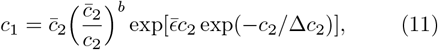

with four parameters: 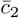 is a scaling factor, and in terms of the tie-line vector in the concentration space *κ*_*α*_ = *k*_*α*_*/N*_*α*_*v*_0_, the reentrant parameters [Fig. 2(b)] are

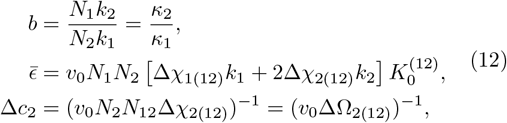

where *b* encodes the phase separation propensity in terms of local tie-line gradient, 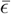 quantifies the effective threebody repulsion and thus reentrant dissolution, and Δ*c*_2_ sets the decay correction scale of effective three-body repulsion. The functional form of Eq. (11) predicts that the concentration scale 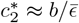 marks the crossover from the phase separation regime to the reentrant dissolution regime. This corresponds, in the free energy Eq. (10), to the three-body terms becoming comparable to the two-body term as *ϕ*_2_ increases.

The calculation can be generalized to *M* component system with *M* (*M* + 1)*/*2 distinct dimers [Eq. (B1)], where we show that the emergent three-body repulsion is a general consequence of integrating out dimer states (Appendix B). We therefore define a general competition 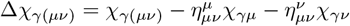 and a totally symmetric three-body interaction 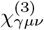 .

## III. PHYSICAL INTERPRETATIONS

### A. Microscopic explanation to model parameters

Here we discuss the physical interpretation of the parameters and how they control the phase boundary geometry in Fig. 2(b). At low concentrations, two-body attraction *χ*_*αβ*_ *<* 0 drives phase separation against entropic mixing, described by the power-law (*c*_2_)^*−b*^ [36]. As *c*_2_ increases, the three-body repulsion dominates, driving reentrant dissolution described by a modified exponential scaling exp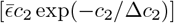. The existence of reentrant dissolution geometry [Fig. 2(b)] requires a positive sign of the competition Δ*χ*. A negative competition Δ*χ*_2(12)_ ∝ Ω_2(12)_ −Ω_12_ *<* 0 rules out the possibility of reentrant dissolution, as dimerization is always favored and increasingly complex polymers are formed, leading to phase-separated condensates. However, for a positive competition Δ*χ*_2(12)_ ∝ Ω_2(12)_ −Ω_12_ *>* 0, monomer–dimer interactions are less favorable than monomer–monomer interactions, producing effective three-body repulsion [Fig. 1(b)-(d)]. Dimerization typically consumes accessible valency, covers surface interaction patches, attenuates electrostatic interactions through charge neutralization and screening, and introduces steric constraints. Experimentally identified mechanisms of reentrant dissolution, such as electrostatic bridging [18], charge inversion [17], and multivalent binding, correspond to the specific microscopic realizations of the interaction competition. In each case, the monomer-dimer channel reduces the ability of the dimer state to participate in further network formation. The concept of competition reveals how effective three-body repulsion can emerge from purely attractive two-body interactions through reversible association and surrounding modulation of binding equilibrium.

The association constant *K*^(12)^ [Eq. (5)] carries concentration dependence to leading order, causing the effective three-body parameter 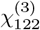 to decay exponentially and vanish at a much higher concentration usually beyond access. This modification is not ad hoc but required for thermodynamic consistency: if the free energy includes a competition-induced effective three-body term, then the mass-action equilibrium conditions [Eqs. (A7)–(A8)] must include the corresponding first-order competition correction. Physically, this reflects the fact that even if two monomers bind strongly in isolation (very negative *ϵ*_12_), dimerization in a crowded interacting environment can be disfavored because the dimer interacts less favorably with its surrounding monomers than the two free monomers it replaces. Moreover, such concentration dependence of the association constant admits higher-order corrections compactly, which is necessary when the approximation Eq. (A13) that requires *K*^(12)^*ϕ* ≪ 1 is not fully met in experiments, suggesting a critical concentration above which a simple exponential scaling [36] is not appropriate. We further investigate in Appendix A 1 and show that implementing the exponential correction 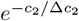 to the scaling 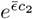 can also capture the depletion corrections to the mass-action equilibrium Eq. (A10) as a mean-field resummation and set the critical concentration scale *c*_2_*/*Δ*c*_2_ ≪1. Above the concentration scale, the effective repulsion [Eqs. (8)–(9)] vanishes. This indicates a regime where the assumptions underlying our effective theory break down, suggesting the emergence of physics not captured by our description. Above this concentration scale, residual two-body attractions (eg. non-ionic [14]) may dominate and re-stabilize phase separation, leading to reentrant condensation. In practice, this regime usually exceeds the physiological and solubility limit. Therefore, a high concentration phase space is rarely accessed in experiments and requires further theoretical investigation.

### B. Phase boundary geometry controlled by the physical parameters

The phase boundary model [Eq. (11)] contains one overall scaling parameter, 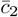 , and three geometry-determining parameters: *b*, 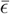 , and Δ*c*_2_. We test how variations in their underlying physical parameters shift the phase boundary as shown in Fig. 3. First, increasing parameter *b* = *κ*_2_*/κ*_1_, corresponding to a larger tie-line gradient [Fig. 3(e)] in concentration space, increases the phase-separated coexistence width in *c*_2_ direction. Therefore, phase separation is delayed [Fig. 3(a)]. The parameter *b* also controls the asymmetry between components in reentrant dissolution. The reentrance in *c*_2_ weakens when *κ*_2_*/κ*_1_ increases, suggesting a relatively broader coexistence region in *c*_2_ direction. Second, 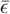 is proportional to the emergent three-body repulsion *χ*^(3)^ such as *χ*_112_ and *χ*_212_. A larger repulsion [Fig. 3(f)] enhances reentrant dissolution. Increasing 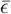 shifts the reentrant boundary to lower concentrations, which represents stronger reentrant dissolution in Fig. 3(b). Third, we examine the competition Δ*χ* [Fig. 3(g)]. Effective three-body repulsion has competing contributions from Δ*χ* of the form *χ*^(3)^ ∝ Δ*χ* exp(−Δ*χc*_2_). However, in reentrant dissolution regime where approximation Eq. (6) is valid, larger competition generally strengthens repulsion, leading to more pronounced reentrance in Fig. 3(c). The exponential suppression term exp(−Δ*χc*_2_), which is stronger when competition Δ*χ* is larger, leads to a less steep reentrant boundary at high *c*_2_. Lastly, we increase the dimer dissociation constant *K*_D_ which is the inverse of the intrinsic association constant 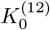 [Fig. 3(h)]. The effective three-body repulsion becomes weaker, and the reentrant dissolution becomes less pronounced [Fig. 3(d)]. Physically, the emergent three-body repulsion arises because dimers interact less favorably with dimers than with monomers. The strength of the effective repulsion mechanism therefore scales with the amount of dimers formed, as described by *K*_D_. In the large *K*_D_ limit where dimerization is suppressed, the effective three-body repulsions [Eqs. (8)–(9)] are negligible, and the reentrant dissolution disappears. These trends allow for the interpretation of experimentally observed shifts of the phase boundaries in terms of the underlying physical parameters, as shown in Secs. IV and Appendix D.

**FIG. 3.**
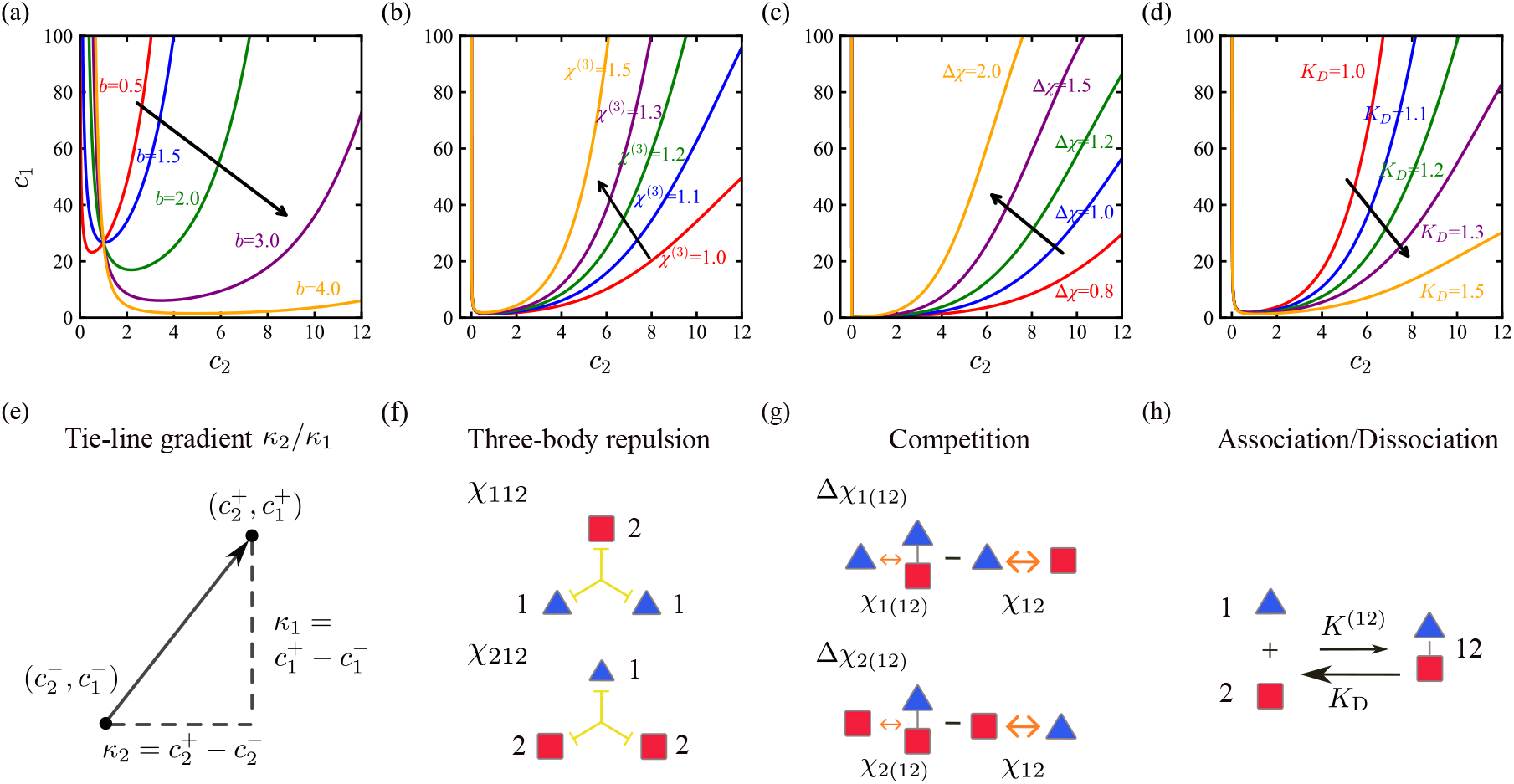
Phase boundary geometry controlled by physical parameters. The region above the curves is in phase-separated state. The corresponding mechanisms are presented schematically below each parameter panel. (a) Increasing *b* = *κ*_2_*/κ*_1_ shifts phase separation to higher *c*_2_. (b) Increasing the effective three-body repulsion *χ*^(3)^ pushes the reentrant dissolution earlier to the lower *c*_2_. (c) Increasing competition Δ*χ* alone pushes reentrant dissolution earlier but the phase boundary less steep at high *c*_2_. (d) Increasing the dimer dissociation constant *K*_D_ delays and weakens the reentrant dissolution by decreasing the effective three-body repulsion. (e) The tie-line gradient 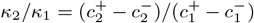 is the orientation of tie-line vector which connects dilute composition 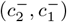 to dense composition 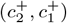 at the two sides of phase-separated region. (f) The effective three-body repulsion *χ*_112_ and *χ*_212_ among three polymers. (g) The interaction competition Δ*χ*_1(12)_ and Δ*χ*_2(12)_ generate the effective three-body repulsion. It is positive when the monomer-dimer channel *χ*_1(12)_ and *χ*_2(12)_ is less attractive than the monomer-monomer channel *χ*_12_. (h) The association constant *K*^(12)^ and dissociation constant *K*_D_ = 1*/K*^(12)^ describe mass-action equilibrium of the dimerization process.

## IV. APPLICATION TO EXPERIMENTAL DATA

To test the theory, we apply our model [Eq. (11)] to the G3BP1 (component-1) - poly(A) RNA (component-2) - suramin (component-3) system, which exhibits reentrant dissolution on 𝒮_12_ plane [Fig. 4(a)]. G3BP1 protein forms large network structure with RNA [39], corresponding to the observed condensate in experiments. This is captured by examining G3BP1-RNA (12) dimer in our model. Suramin is a highly polyanionic small molecule modulator which disrupts G3BP1-RNA binding and suppresses condensation [38]. This modulation effect was studied by measuring the hydrodynamic radius *R*_H_ of G3BP1 at different [RNA] values reported in Ref. [38]. With 0 µM suramin, *R*_H_ increases with [RNA], suggesting G3BP1-RNA binding, from which the G3BP1-RNA *K*_D_ = 127 ng/µl (95% CI: 18 − 253 ng/µl) was inferred. With 10 µM suramin, the increase of *R*_H_ was completely eliminated, which suggests strong suppression of G3BP1-RNA interaction and binding.

**FIG. 4.**
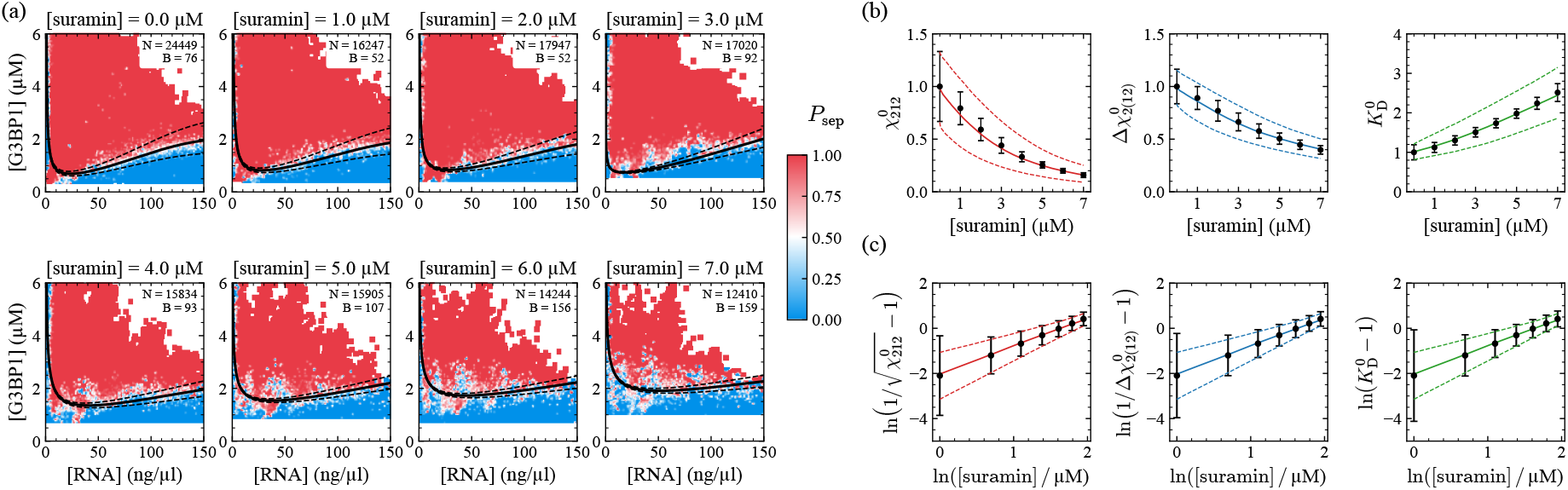
Apply the model to G3BP1-RNA _12_ plane using Eq. (11) with free parameters 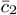 and *b*, and globally constrained parameters 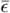 and Δ*c*_2_ (Appendix C). (a) Globally fitted G3BP1-RNA phase boundary of PhaseScan measurement at eight [suramin] = 0.0 µM ∼ 7.0 µM, with covariance error envelopes. The raw data are shown as a heatmap of the phase-separated probability *P*_sep_. The number of measurements *N* and extracted boundary points *B* are recorded for each [suramin] slab. (b) The inferred and normalized effective three-body repulsion 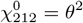 , competition 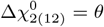 and G3BP1-RNA dissociation constant 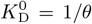 (dot with error bar) are described by Hill function *θ* = *θ*([suramin]) = 1*/*(1 + ([suramin]*/K*_sur_)^*n*^) with (c) Plotting the Hill-transformed quantities ln 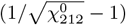 , ln 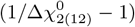 , and ln 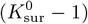 versus ln[suramin] confirms inferred *K*_sur_ and *n* (solid curve). Envelope (dashed curve) represents ±1*σ* uncertainty in the normalized Hill prediction. linearity with slope *n* and y-intercept *−n* ln *K*_D_ [Eq. (C1)]. [suramin] = 0.0 µM point is excluded due to logarithmic divergence under this transform.

PhaseScan [40] is a high-throughput platform for measuring multicomponent phase diagrams. PhaseScan measurements [38] show that the reentrant phase boundary becomes less steep and the reentrant dissolution becomes weaker on the G3BP1–RNA 𝒮_12_ plane as the suramin concentration increases [Fig. 4(a)]. These observations imply that the reentrant geometry parameters, and therefore the underlying physical parameters such as effective three-body repulsion *χ*_212_, competition Δ*χ*_2(12)_, and dissociation constant *K*_D_, must explicitly depend on the concentration of suramin. The corresponding interaction and association between the G3BP1 and RNA scale with the probability that their binding is not disrupted by suramin, which is quantitatively described by the Hill function

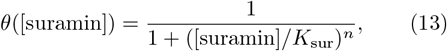

where *K*_sur_ is the suramin dissociation constant, and *n* is the Hill coefficient that quantifies cooperative binding. More importantly, the Hill-function fit is intrinsically scale-invariant and therefore does not require determination of the absolute magnitudes of the physical parameters. We thus performed constrained global fits across all [suramin] concentrations for 𝒮_12_ plane, imposing the Hill function dependence (see Appendix C), which reduced the total number of fitting parameters and tested the internal consistency of our microscopic framework.

We investigate the phase boundary on the G3BP1 - RNA 𝒮_12_ plane at eight suramin concentrations from 0.0µM to 7.0 µM. To capture the microscopic modulation mechanism of suramin, we restrict the competition 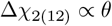 ([suramin]), *K*_D_ ∝ 1*/θ*([suramin]), and hence 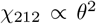 ([suramin]). *θ*([suramin]) is the Hill function defined in Eq. (13) with parameters *K*_sur_, the suramin dissociation constant, and *n*, the suramin binding Hill coefficient. The Hill function serves as a minimal statistical model to capture the suramin modulation effect in terms of competitive binding. The phase boundary model is Eq. (11), and here it reads:

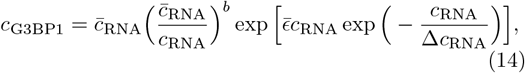

which captures the reentrant boundaries in Fig. 4(a). The inferred parameters *χ*_212_ and Δ*χ*_2(12)_ decrease, while *K*_D_ increases with [suramin]. The functional dependence on [suramin], shown in [Fig. 4(b)–(c)], is captured by the Hill model with *K*_sur_ = 5.08 ± 1.49 µM and *n* = 1.29 ± 0.47. The inferred *K*_sur_ 5 ≈ µM lies midway between [suramin] = 0.0 µM, where G3BP1-RNA interaction is intact, and [suramin] = 10.0 µM, where G3BP1-RNA interaction is fully disrupted [38]. The Hill coefficient *n* ≈ 1 suggests that suramin binding has no cooperativity. Because *K*_sur_ and *n* extracted from Hill constraint are scale-free, uncertainties in *N*_*α*_ and *v*_0_ primarily rescale absolute *χ*_212_, Δ*χ*_2(12)_, and *K*_D_ but do not affect the inferred Hill parameters.

Lastly, we examine whether our approximation [Eq. (6)] is valid and self-consistent in the global fitting. Using the estimates in Appendix C, we show that the simple exponential scaling [Eq. (7)] breaks down at approximately 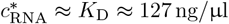 , which captures the beginning part of reentrant dissolution boundary but indicates the necessity of the modified scaling form *cxe*^*−px*^ beyond this concentration. By contrast, the perturbative validity condition for the modulated exponential [Eq. (6)] extends to a broader concentration regime, which, from the global fitting (Δ*c*_RNA_)^*−*1^ *<* 0.0043 (ng/µl)^*−*1^ [41], gives 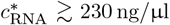. With increasing [suramin], the perturbative condition is increasingly well satisfied because of the decrease in Δ*c*_RNA_. This supports the internal consistency of the theory and its application to experimentally relevant concentrations.

## V. DISCUSSION

Our model is formulated within a coarse-grained meanfield framework. However, the emergence of an effective three-body interaction is not specific to the Flory– Huggins lattice. The key is to eliminate the reversibly formed dimer coupled with interaction-modulated mass-action equilibrium. Integrating out this dimer degree of freedom generates higher-order terms proportional to the product of interaction-channel competition and the association constant. This structure and mechanism apply equally to continuum density expansions and other mean-field free-energy formulations, while the lattice description serves only as a transparent analytical vehicle rather than a fundamental requirement.

In the perturbative calculation, we expand the free energy to third order in volume fraction under dilute conditions and assume that the effective association constant varies weakly with concentration at leading order. These approximations provide controlled analytic access to reentrant behavior and are appropriate for the experimental PhaseScan measurements analyzed here. At high concentrations, additional higher-order contributions are expected.

The phase boundary equation contains free parameters that are decoupled across different concentration regimes. We approximate the tie-line *k*_*α*_ as piecewise constant within each of the phase separation and reentrant dissolution regimes, while allowing the tie-line to differ across the regimes. Hence, the tie-line *k*_*α*_ in each parameter [Eq. (12)] is better interpreted as the local tie-line vector. This is supported by the observation that the boundary curvature remains approximately uniform within each regime [Figs. 4(a) and 5(a)].

## ACKNOWLEDGMENTS

This work has received support from Transition Bio Limited. (D.Q. and R.M.S.). The authors thank Ella de Csilléry for many discussions.

The authors declare no conflict of interest.

## DATA AVAILABILITY

The data and analysis scripts that support the findings of this article are openly available. [41]

## Appendix A: Two-component

### 1. Derivation for free energy with many-body interactions

The two-component Flory–Huggins free energy is

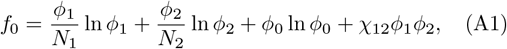

where we set *k*_*B*_*T* = 1 for simplicity. In the dimerization process 1 + 2 ↔ 12 in the homogeneous state, we introduce a dimer component 12 explicitly labeled by its constituent monomers, with a free energy of formation *ϵ*_12_. When the dimer participates in interactions, we enclose it in parentheses, (12), to indicate that it is treated as a single independent species. The Flory–Huggins free energy after dimerization is

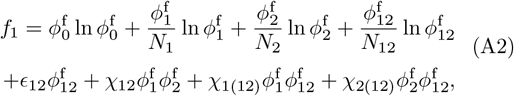

where the superscript f denotes the volume fraction of the free monomers which are not in the dimer after dimerization. To integrate out the dimer state 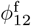, first consider the conservation of the number of monomer units in terms of volume fraction

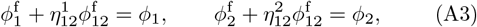

where we define a volume factor 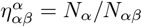 , the frac-tion of monomer *α* in dimer *αβ*. We assume the incompressibility condition *N*_12_ = *N*_1_ + *N*_2_, so 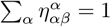. At equilibrium, *f*_1_ is minimized under this conservation con-straint with respect to 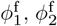 , and 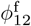 , using Lagrangian multipliers *µ*_1_ and *µ*_2_

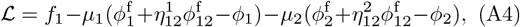

which leads to chemical potentials 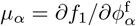

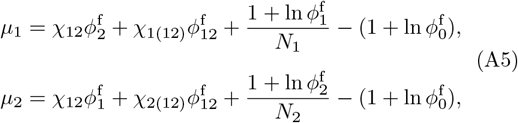

and chemical potential equilibrium for dimer 12

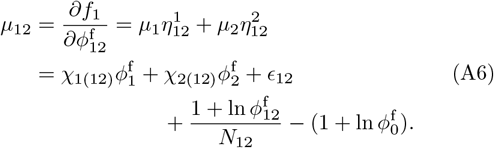

Chemical potentials explicitly include the contribution of the many-body interaction 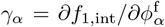 as a func-tion of 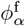. Substituting Eq. (A5) into Eq. (A6) which is the chemical potential equilibrium, canceling 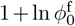 on both sides and shifting *ϵ*_12_ − 1*/N*_12_ → *ϵ*_12_, we have

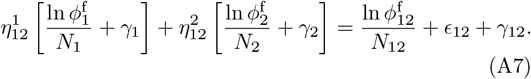

Multiplying by *N*_12_ on both sides, we obtain

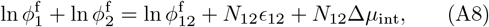

where Δ*µ*_int_ collects the many-body interaction contributions. To the leading order, the interaction contribution

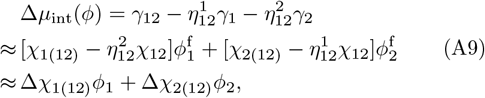

where the competition parameters Δ*χ*_1(12)_ and Δ*χ*_2(12)_ appear as the linear coefficients.

To integrate out the dimer state, we solve for 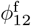 from the iterative equation based on the conservation condition Eq. (A3) self-consistently

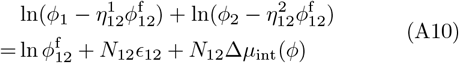

First, examine the zeroth-order solution where we ap-proximate 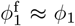 and 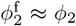. This approximation is controlled when

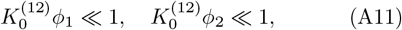

which would lead to an exponential scaling law is valid [36]. In the absence of competition Δ*µ*_int_ = 0, we identify the intrinsic association constant independent of *ϕ* with

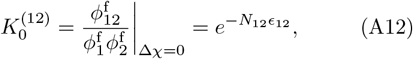

where we identify the Gibbs free energy change Δ*G*_0_ with *N*_12_*ϵ*_12_ through Δ*G*_0_ = −ln *K*_0_. However, we switch on the interaction and propose an effective association constant with a weak *ϕ* dependence

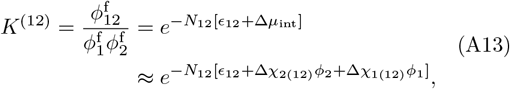

For experimentally relevant parameter regimes, the condition Eq. (A11) may not be well satisfied, where we performed extra phenomenological fitting as in Ref. [36]. This motivates us to solve the iterative equation Eq. (A10) for 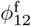 to higher-order and include the depletion correction when 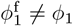 and 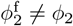

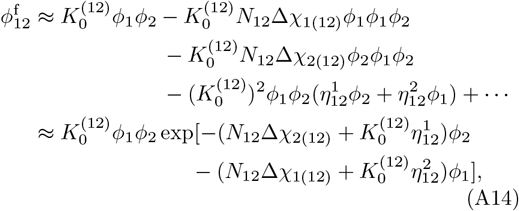

where the first line gives the leading correction obtained by one iteration of the nonlinear mass-action equation Eq. (A10), while the exponential form is a low-density resummation of these power-series depletion corrections. In this representation, competition and depletion generate the same correction functional form to the mass-action equilibrium. For reentrant systems, the depletion contribution 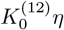 and the interaction competition contribution *N*_12_Δ*χ* naturally enter at comparable parametric order. This is consistent with the emergent effective threebody repulsion 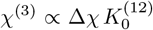 , which relates the competition coefficient to the underlying binding strength. The depletion correction may therefore be absorbed phenomenologically into an effective competition parameter. This operation extends our regime of validity to a larger concentration range, provided

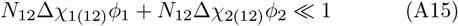

We calculate the perturbation of the interaction free energy Δ*f*_int_ = *f*_1,int_ − *f*_0,int_ term by term using the constraint [Eq. (A3)]

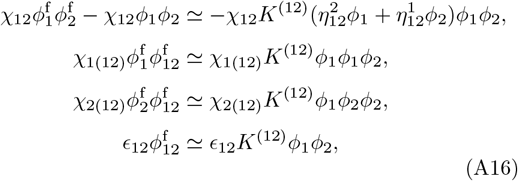

where the previous two-body interaction terms generate three-body terms, and the 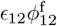 term contributes to the two-body term. Define the effective three-body interaction parameters with weak concentration dependence

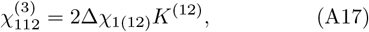

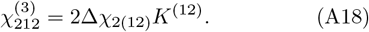

We thus obtain the renormalized free energy *f* ≈ *f*_0_ + Δ*f*_int_ with emergent three-body interactions [Eq. (10)]. A fully consistent saddle-point elimination also includes consideration of the translational entropy of the dimerization perturbation Δ*f*_entropy_

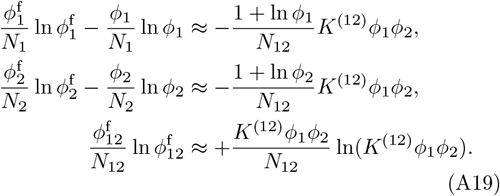

Collecting results, we find that the terms in the form *ϕ*^2^ ln *ϕ* completely cancel, leaving only a two-body renormalization

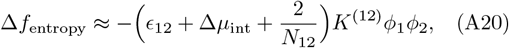

while the two-body interaction contribution is

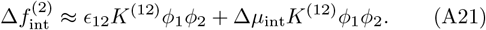

Therefore, the two-body terms take the form

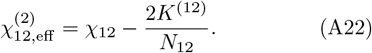

The phase-boundary analysis below depends only on the three-body terms, this two-body renormalization does not affect the mathematical structure of the theory, but the *ϕ* dependence generates contribution to three-body terms. We expand

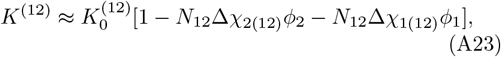

which generates an additional three-body term

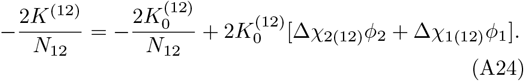

The expansion generates only 𝒪 (*ϕ*^4^) terms compared to the previous three-body terms

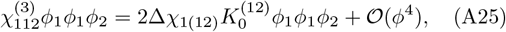

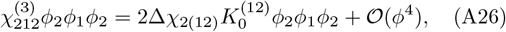

Collecting results, in the constant-coefficient picture, the three-body interaction is only rescaled

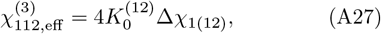

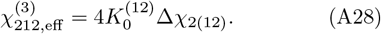

We will work with the intuitive and compact form of free energy with *ϕ* dependent coefficients Eq. (A17)–(A17), and derive the phase boundary equation from Eq. (10).

#### B. Analytical phase boundary via tie-line analysis

Tie-line analysis [35] provides a geometric method for calculating the phase boundary normal vector *n*_*α*_, which has contributions from solute entropy *Q*_*α*_, solvent entropy *Q*_0_ (usually ignored as *ϕ*^0^ ≈ 1), and many-body interactions *X*_*α*_ adapted from Ref. [36]. 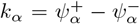 is the *α* component of the tie-line vector [Fig. 2]. The phase boundary normal vector *n*_*α*_ can be written as an expansion around the dilute phase volume fraction 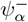

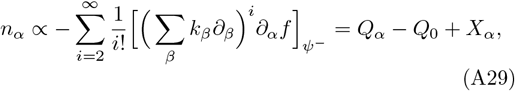

where *f* = *f*_entropy_ + *f*_int_ is the free energy. The translational entropy is approximated as

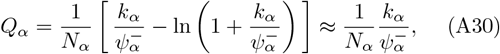

where we assume 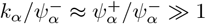. The three-body interaction contribution has the general form

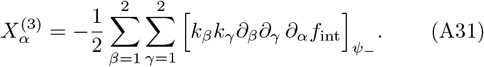

The systems are usually reentrant only in 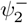 direction in probing limit, so we work in strong enrichment regime in 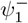. The normal vector is thus 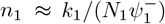. To leading order in the probing-limit, we therefore retain only the 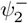 dependence of 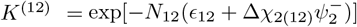. From Eq. (10) with shorthand *a* = *N*_12_Δ*χ*_2(12)_, we calculate normal vector *n*_2_

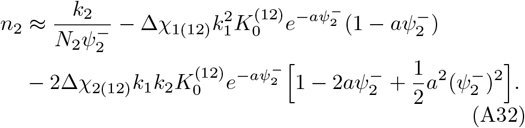

The first interaction term integrates as ∫ (1−*ax*)*e*^−*ax*^*dx* = *xe*^−*ax*^ + *C* while the second as 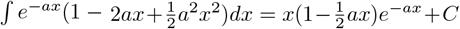. Therefore, solving the geometric equation 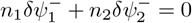 gives

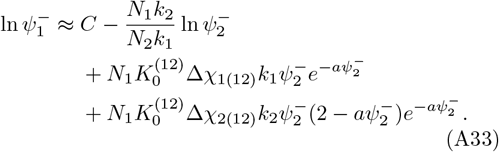

In the controlled validity regime Eq. (A15), which implies 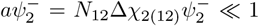, the exact second contribution differs from 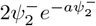 only by a higher-order correction 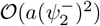. We therefore obtain

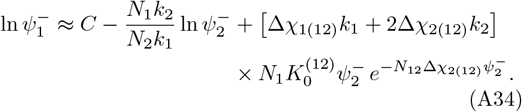

where *C* is an integration constant. Rewrite in terms of molar concentrations using 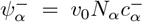 , the model has functional form [Eq. (11)] with four parameters [Eq. (12)].

## Appendix B: M-component

We generalize the calculation in Appendix A to *M* - component labeled as 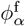, *α* = 1, 2, …, *M*. We include *M* (*M* + 1)*/*2 dimer species formed from pairs of monomers, denoted by 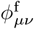 , 1 ≤ *µ* ≤ *ν* ≤ *M*. These include *M* self-dimers 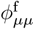 which describe homotypic phase separation. A free energy penalty *ϵ*_*µν*_ = *ϵ*_*νµ*_ is assigned to form a dimer component 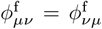. The two-body interaction matrix *χ*_*αβ*_ is symmetric *χ*_*αβ*_ = *χ*_*βα*_ with vanishing diagonal entries *χ*_*αα*_ = 0. The free energy is

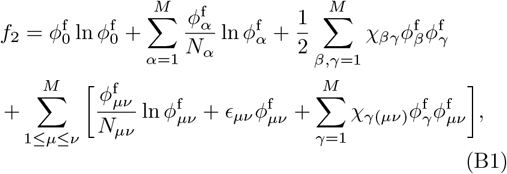

with solvent volume fraction 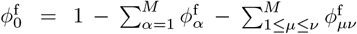. We define stoichiometric coefficients 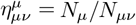 with symmetry 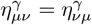. There are *M* volume conservation constraints 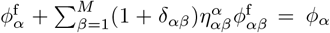. Minimizing free energy, we obtain the chemical potential of the monomer *α*

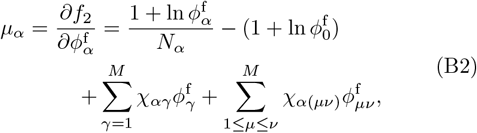

and the chemical potential of the dimer component *µν*

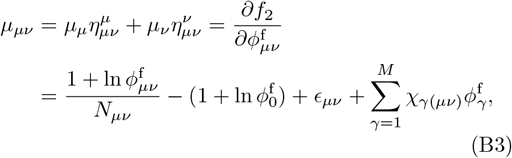

Define the multicomponent competition

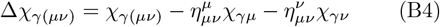

and equivalently the molecular competition ΔΩ_*γ*(*µν*)_ = *N*_*γ*_*N*_*µν*_Δ*χ*_*γ*(*µν*)_ = Ω_*γ*(*µν*)_ −Ω_*γµ*_ −Ω_*γν*_. To first order in *ϕ*^*α*^, the effective association constant derived from equilibrium condition [Eq. (B3)] is

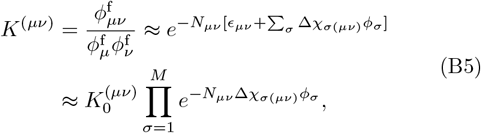

where each relevant competition 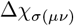 contributes to disfavoring dimerization. The three-body 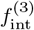 includes the monomer-dimer interaction channel term

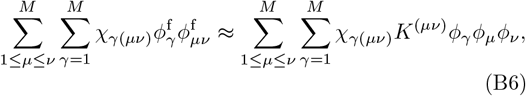

and the monomer-monomer channel term

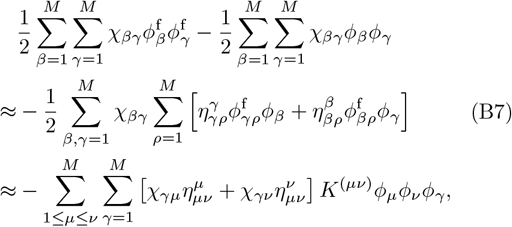

where we use index symmetry in 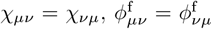 and 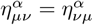, and relabel dummy indices at the second equality. Adding these two terms reproduces the competition 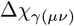. The renormalized three-body interaction free energy is

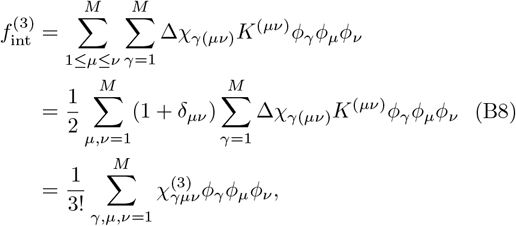

where the factor 1 + *δ*_*µν*_ corrects the counting for self-dimers. The totally symmetric three-body interaction parameter is defined as

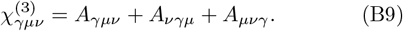

We impose symmetry in dimer indices, *A*_*γµν*_ = *A*_*γνµ*_. To obtain the correct prefactor, *A*_*γµν*_ is nonzero only when the dimer *µν* is physically present in the system, and is defined as

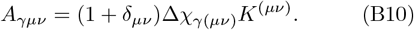

The normal vector has an interaction contribution

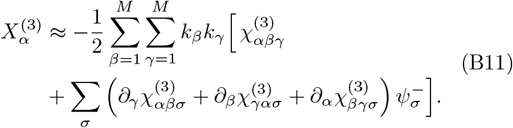

To capture common experimental observations and maintain analyticity, we assume that only component *α* enters the exponential modulation for reentrant dissolution along 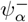. Therefore, *n*_*α*_ only has 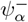 dependence.

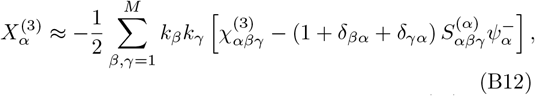

where the correction term due to decay in *K*^(*µν*)^ is

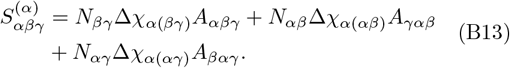

Phase boundary equation 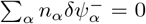 can be solved analytically with appropriate approximation.

## Appendix C: Global fit details

We infer the effective physical parameters under the lattice-volume and tie-line assumptions: the competition Δ*χ*_2(12)_ = (*v*_0_*N*_2_*N*_12_Δ*c*_2_)^*−*1^, the concentration-independent three-body repulsion 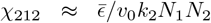, and the intrinsic dissociation constant 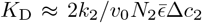. Such inference requires estimating the polymer length *N*_*α*_, the referencing lattice volume *v*_0_ = 18 g/mol ÷10^3^ g/L = 1.8 × 10^*−*2^M^*−*1^ corresponding to a water molecule, and a typical tie-line magnitude 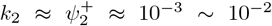. This follows from that the typical dense phase volume fraction 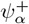 is much larger than the dilute phase volume fraction 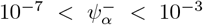. The following estimates are made: G3BP1, an RNA-binding protein with 466 amino acids and molecular weight 52 kDa [4], has contour length (scales with number of monomer units) *N*_1_ = 5.2× 10^4^ Da÷ 18 Da = 3 × 10^3^ compared with water (18 Da). The linear chain poly(A)-RNA (700− 3500kDa) has typical contour length *N*_2_ = 2.1× 10^3^ kDa ÷18 Da ≈10^4^. An average value 2100 kDa per RNA molecule is used to convert 1ng/µl = 10^*−*3^g/l = 0.48nM. Suramin has typical length *N*_3_ = 1.3 kD ÷ 18 Da≈10^2^. These estimates provide a crude inference of physical parameters, though the global fitting results are scale invariant.

We performed global fitting to G3BP1-RNA 𝒮_12_ subspace at eight different [suramin] slices from PhaseScan data [Fig. 4(a)]. Each piece of the phase boundary requires four parameters in our model, so we have 4 × 8 = 32 parameters. However, the two reentrant parameters 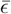 and Δ*c*_2_ have specific [suramin] dependence pinned by Hill function describing interruption from suramin binding. To be specific, 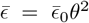 and 1*/*Δ*c*_2_ = (1*/*Δ*c*_2,0_) × *θ* are constrained by the Hill function *θ* = *θ*([suramin]) = 1*/*(1 +([suramin]*/K*_sur_)^*n*^), up to proportionality constants 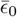 and Δ*c*_2,0_. The Hill function only has two parameters, so the total number of fitting parameters reduces to 2 × 8 + 2 = 18 for global fit.

Phase boundary points were extracted from each measured PhaseScan slice, where the data points were classified in terms of phase separation probability as either in the homogeneous state (*P*_sep_ = 0) or in the phase-separated state (*P*_sep_ = 1). The phase boundary was extracted by estimating the conditional probability of phase separation using binned histograms, Gaussian , smoothing, and identifying the contour where *P*_sep_ ≈0.5. The high-throughput measurement provides 𝒪 (10^4^) data points on each phase diagram, among which 𝒪 (10^2^) are extracted as phase boundary points. Fitting parameters were obtained by bounded nonlinear least-squares minimization (curve_fit function in SciPy optimization library). Parameter uncertainties were estimated from the covariance matrix. The global optimization for G3BP1-RNA converged after 36 function evaluations [41].

To evaluate the quality of Hill fits in the context of the global fits, we linearize the functional form and examine deviations from straight-line behavior. In general, for a quantity 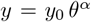 where *θ*[L] = 1*/*(1 + ([L]*/K*_L_)^*n*^), we apply the standard Hill transformation

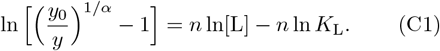

Hence, plotting ln[(*y*_0_*/y*)^1*/α*^ − 1] versus ln[L] gives a straight line with slope *n* and intercept −*n* ln *K*_L_.

## Appendix D: AMP-RNA validation

To examine the generality of the model, we also investigate anti-microbial peptides (AMP, component 1) and Poly(A)-RNA (component 2) system [42]. AMPs are short, highly cationic peptides, while RNA is highly polyanionic. AMP-RNA binding and multivalent electrostatic attraction lead to complex formation and liquid-liquid phase separation [43]. We therefore consider the AMP-RNA dimerization process. The condensate dis-solves when RNA concentration is further increased [42]. Take AMP *N*_1_≈ 1 kDa ÷18 Da≈ 10, RNA *N*_2_ ≈10^4^,and thus *N*_12_ = *N*_1_ + *N*_2_ ≈10^4^. Experiment [42] indicates similar phase separation behavior across the three AMPs. We therefore fix a global scaling parameter 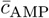 and a global phase separation parameter *b* for all three AMPs. The phase boundary fitting based on Eq. (11) and the result shown in Fig. 5(a) suggest a consistent order in concentration-independent three-body repulsion *χ*_212_, competition Δ*χ*_2(12)_ and AMP-RNA dissociation constant *K*_D_ in Fig. 5(b). Inferred three-body repulsion *χ*_212_ = 2Δ*χ*_2(12)_*/K*_D_ follows the order: Buforin-2 *>* Os-C *>* P113. This trend is consistent with experimental observations [Fig. 5(a)] that Buforin-2 exhibits the steepest reentrant dissolution boundary among the AMPs as a consequence of strong AMP–RNA binding and pronounced charge-neutralization–driven self-limiting behavior at high RNA concentrations [36, 42]. In contrast, P113 shows the strongest intrinsic phase-separation tendency and therefore the latest onset of reentrant dissolution [42]. The fitted parameters also show an increasing trend in competition Δ*χ*_2(12)_ and dissociation constant *K*_D_, although Os-C and Buforin-2 have comparable *K*_D_. Experiments indicate that the RNA-AMP binding affinity follows the order: P113 *>* Os-C *>* Buforin-2, corresponding to 1*/K*_D_ which is qualitatively captured by our inference [42].

**FIG. 5.**
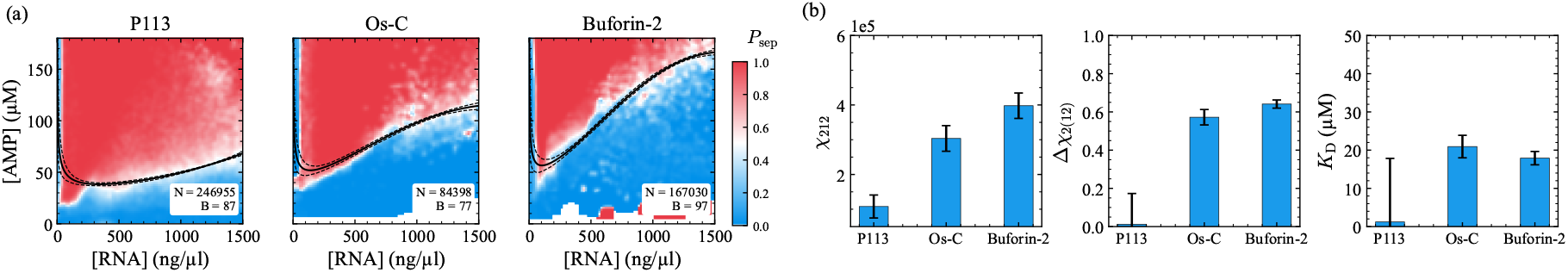
Experimental validation on AMP-RNA𝒮 _12_ plane. (a) Phase boundaries of three AMPs (P113, Os-C, Buforin-2) against [RNA] are fitted with the model Eq. (11). The raw data are shown as a heatmap of the phase separation probability *P*_sep_. The number of measurement *N* and extracted boundary points *B* are recorded for each AMP panel. (b) The inferred three-body repulsion *χ*_212_, competitionΔ*χ*_2(12)_, and AMP-RNA dissociation constant *K*_D_ are plotted in histograms, where we observe increasing order in the three physical parameters.

## References

[1] N. A. Erkamp, T. Sneideris, H. Ausserwöger, D. Qian, S. Qamar, J. Nixon-Abell, P. St George-Hyslop, J. D. Schmit, D. A. Weitz, and T. P. Knowles, Spatially non-uniform condensates emerge from dynamically arrested phase separation, Nat. Commun. 14, 684 (2023).

[2] P. R. Banerjee, A. N. Milin, M. M. Moosa, P. L. Onuchic, and A. A. Deniz, Reentrant phase transition drives dynamic substructure formation in ribonucleoprotein droplets, Angew. Chem. Int. Ed. 56, 11354 (2017).

[3] S. Boeynaems, S. Alberti, N. L. Fawzi, T. Mittag, M. Polymenidou, F. Rousseau, J. Schymkowitz, J. Shorter, B. Wolozin, L. Van Den Bosch, et al., Protein phase separation: a new phase in cell biology, Trends Cell Biol. 28, 420 (2018).

[4] P. Yang, C. Mathieu, R.-M. Kolaitis, P. Zhang, J. Messing, U. Yurtsever, Z. Yang, J. Wu, Y. Li, Q. Pan, et al., G3BP1 is a tunable switch that triggers phase separation to assemble stress granules, Cell 181, 325 (2020).

[5] A. Carlson and L. Mahadevan, Elastohydrodynamics and kinetics of protein patterning in the immunological synapse, PLoS Comput. Biol. 11, e1004481 (2015).

[6] Y. Shin and C. P. Brangwynne, Liquid phase condensation in cell physiology and disease, Science 357, eaaf4382 (2017).

[7] A. W. Folkmann, A. Putnam, C. F. Lee, and G. Seydoux, Regulation of biomolecular condensates by interfacial protein clusters, Science 373, 1218 (2021).

[8] S. Alberti and A. A. Hyman, Biomolecular condensates at the nexus of cellular stress, protein aggregation disease and ageing, Nat. Rev. Mol. Cell Biol. 22, 196 (2021).

[9] T. C. Michaels, A. Šarić, S. Curk, K. Bernfur, P. Arosio, G. Meisl, A. J. Dear, S. I. Cohen, C. M. Dobson, M. Vendruscolo, et al., Dynamics of oligomer populations formed during the aggregation of alzheimer’s aβ42 peptide, Nat. Chem. 12, 445 (2020).

[10] T. C. Michaels, D. Qian, A. Šarić, M. Vendruscolo, S. Linse, and T. P. Knowles, Amyloid formation as a protein phase transition, Nat. Rev. Phys. 5, 379 (2023).

[11] C. P. Brangwynne, P. Tompa, and R. V. Pappu, Polymer physics of intracellular phase transitions, Nat. Phys. 11, 899 (2015).

[12] J.-M. Choi, A. S. Holehouse, and R. V. Pappu, Physical principles underlying the complex biology of intracellular phase transitions, Annu. Rev. Biophys. 49, 107 (2020).

[13] M. T. Guo, A. Rotem, J. A. Heyman, and D. A. Weitz, Droplet microfluidics for high-throughput biological assays, Lab Chip 12, 2146 (2012).

[14] G. Krainer, T. J. Welsh, J. A. Joseph, J. R. Espinosa, S. Wittmann, E. de Csilléry, A. Sridhar, Z. Toprak-cioglu, M. Gudič, R. Collepardo-Guevara, and T. P. J. Knowles, Reentrant liquid condensate phase of proteins is stabilized by hydrophobic and non-ionic interactions, Nat. Commun. 12, 1085 (2021).

[15] H. Ausserwöger, E. de Csilléry, D. Qian, G. Krainer, T. J. Welsh, T. Sneideris, T. M. Franzmann, S. Qamar, N. A. Erkamp, J. Nixon-Abell, et al., Quantifying collective interactions in biomolecular phase separation, Nat. Commun. 16, 7724 (2025).

[16] S. F. Banani, H. O. Lee, A. A. Hyman, and M. K. Rosen, Biomolecular condensates: organizers of cellular biochemistry, Nat. Rev. Mol. Cell Biol. 18, 285 (2017).

[17] I. Alshareedah, T. Kaur, J. Ngo, H. Seppala, L.-A. Djomnang Kounatse, W. Wang, M. M. Moosa, and P. R. Banerjee, Interplay between short-range attraction and long-range repulsion controls reentrant liquid condensation of ribonucleoprotein–rna complexes, J. Am. Chem. Soc. 141, 14593 (2019).

[18] Y.-H. Lin, T. H. Kim, S. Das, T. Pal, J. Wessén, A. K. Rangadurai, L. E. Kay, J. D. Forman-Kay, and H. S. Chan, Electrostatics of salt-dependent reentrant phase behaviors highlights diverse roles of atp in biomolecular condensates, eLife 13, RP100284 (2025).

[19] A. N. Semenov and M. Rubinstein, Thermoreversible gelation in solutions of associative polymers. 1. statics, Macromolecules 31, 1373 (1998).

[20] J. Wang, J.-M. Choi, A. S. Holehouse, H. O. Lee, X. Zhang, M. Jahnel, S. Maharana, R. Lemaitre, A. Pozniakovsky, D. Drechsel, et al., A molecular grammar governing the driving forces for phase separation of prion-like rna binding proteins, Cell 174, 688 (2018).

[21] Y.-H. Lin, J. Wessén, T. Pal, S. Das, and H. S. Chan, in Phase-Separated Biomolecular Condensates: Methods and Protocols (Springer, 2022) pp. 51–94.

[22] M. Farag, A. S. Holehouse, X. Zeng, and R. V. Pappu, Fireball: A tool to fit protein phase diagrams based on mean-field theories for polymer solutions, Biophys. J. 122, 2396 (2023).

[23] J. T. Overbeek and M. J. Voorn, Phase separation in polyelectrolyte solutions. theory of complex coacervation, J. Cell. Comp. Physiol. 49, 7 (1957).

[24] Y.-H. Lin, J. Song, J. D. Forman-Kay, and H. S. Chan, Random-phase-approximation theory for sequence-dependent, biologically functional liquid-liquid phase separation of intrinsically disordered proteins, J. Mol. Liq. 228, 176 (2017).

[25] J. McCarty, K. T. Delaney, S. P. Danielsen, G. H. Fredrickson, and J.-E. Shea, Complete phase diagram for liquid–liquid phase separation of intrinsically disordered proteins, J. Phys. Chem. Lett. 10, 1644 (2019).

[26] T. K. Lytle and C. E. Sing, Transfer matrix theory of polymer complex coacervation, Soft Matter 13, 7001 (2017).

[27] T. K. Lytle, L.-W. Chang, N. Markiewicz, S. L. Perry, and C. E. Sing, Designing electrostatic interactions via polyelectrolyte monomer sequence, ACS Cent. Sci. 5, 709 (2019).

[28] J. A. Joseph, A. Reinhardt, A. Aguirre, P. Y. Chew, K. O. Russell, J. R. Espinosa, A. Garaizar, and R. Collepardo-Guevara, Physics-driven coarse-grained model for biomolecular phase separation with near-quantitative accuracy, Nat. Comput. Sci. 1, 732 (2021).

[29] K. L. Saar, A. S. Morgunov, R. Qi, W. E. Arter, G. Krainer, A. A. Lee, and T. P. Knowles, Learning the molecular grammar of protein condensates from sequence determinants and embeddings, Proc. Natl. Acad. Sci. U.S.A. 118, e2019053118 (2021).

[30] P. J. Flory, Thermodynamics of high polymer solutions, J. Chem. Phys. 10, 51 (1942).

[31] M. L. Huggins, Solutions of long chain compounds, J. Chem. Phys. 9, 440 (1941).

[32] C. Luo, Y. Qiang, and D. Zwicker, Beyond pairwise: Higher-order physical interactions affect phase separation in multicomponent liquids, Phys. Rev. Res. 6, 033002 (2024).

[33] A. Chattaraj, M. L. Blinov, and L. M. Loew, The solubility product extends the buffering concept to heterotypic biomolecular condensates, elife 10, e67176 (2021).

[34] S. A. Akram, A. Chattaraj, T. Salava Jr, J. A. Ditlev, L. M. Loew, and J. D. Schmit, Biomolecular phase boundaries are described by a solubility product that accounts for variable stoichiometry and soluble oligomers, J. Am. Chem. Soc. 147, 45471 (2025).

[35] D. Qian, T. J. Welsh, N. A. Erkamp, S. Qamar, J. Nixon-Abell, G. Krainer, P. St. George-Hyslop, T. C. Michaels, and T. P. Knowles, Tie-line analysis reveals interactions driving heteromolecular condensate formation, Phys. Rev. X 12, 041038 (2022).

[36] D. Qian, J. Acker, R. M. Scrutton, C. M. Fischer, S. Qamar, T. Sneideris, A. Borodavka, and T. P. J. Knowles, Scaling relations of multicomponent phase coexistence boundaries, PRX Life 4, 013026 (2026).

[37] D. Qian, H. Ausserwoger, W. E. Arter, R. M. Scrutton, T. J. Welsh, T. Kartanas, N. Ermann, S. Qamar, C. M. Fischer, T. Sneideris, et al., Molecular mechanisms of condensate modulation from energy-dominance analysis, Phys. Rev. Appl. 23, 064017 (2025).

[38] D. Qian, H. Ausserwoger, W. E. Arter, R. M. Scrutton, T. J. Welsh, T. Kartanas, N. Ermann, S. Qamar, C. Fischer, T. Sneideris, et al., Linking modulation of bio-molecular phase behaviour with collective interactions, BioRxiv , 2023 (2023).

[39] J. Guillén-Boixet, A. Kopach, A. S. Holehouse, S. Wittmann, M. Jahnel, R. Schlüßler, K. Kim, I. R. Trussina, J. Wang, D. Mateju, et al., RNA-induced con-formational switching and clustering of G3BP drive stress granule assembly by condensation, Cell 181, 346 (2020).

[40] W. E. Arter, R. Qi, N. A. Erkamp, G. Krainer, K. Didi, T. J. Welsh, J. Acker, J. Nixon-Abell, S. Qamar, J. Guillén-Boixet, et al., Biomolecular condensate phase diagrams with a combinatorial microdroplet platform, Nat. Commun. 13, 7845 (2022).

[41] J. Qiao, Reentrant phase boundary analysis code (2026).

[42] T. Sneideris, N. A. Erkamp, H. Ausserwöger, K. L. Saar, T. J. Welsh, D. Qian, K. Katsuya-Gaviria, M. L. Johncock, G. Krainer, A. Borodavka, et al., Targeting nucleic acid phase transitions as a mechanism of action for antimicrobial peptides, Nat. Commun. 14, 7170 (2023).

[43] S. Maharana, J. Wang, D. K. Papadopoulos, D. Richter, A. Pozniakovsky, I. Poser, M. Bickle, S. Rizk, J. Guillén-Boixet, T. M. Franzmann, et al., RNA buffers the phase separation behavior of prion-like RNA binding proteins, Science 360, 918 (2018).

